# Optimizing Transfection Efficiency in Primary Human Splenic Fibroblast Cells

**DOI:** 10.1101/2025.01.19.633792

**Authors:** Ashpreet Kaur, Victoria A. Assogba, Jovy M. G. Assogba, Barnabe D. Assogba

**Affiliations:** Faculty of Science, Department of Biology & Health Science, Kwantlen Polytechnic University, 12666 72 Avenue, Surrey, BC V3W 2M8 (Canada); Welsh Hill School, 2610 Newark Granville Rd, Granville, OH 43023, United States

**Author notes:** **Corresponding author:** Dr. Barnabe Dossou Assogba (principal author), Faculty of Science, Department of Biology, Kwantlen Polytechnic University, 12666 72 Ave. Surrey, British Columbia, V3W 2M8.

## Abstract

Transfection, a fundamental molecular biology technique, is crucial in introducing foreign DNA into cells. Despite its pivotal importance, mastering this technique remains a formidable challenge. In this study, we employed the calcium phosphate reagent and conducted a systematic exploration through time-dependent experiments to assess its impact on human primary splenic fibroblast cells (HSFC). The results precisely delineate optimal time frames for achieving enhanced transfection efficiency, offering conclusive evidence for the efficacy of calcium phosphate in transient transfection within the cells. The data significantly advances our comprehension of HSFC reactions, contributing to refining transfection methodologies with broader implications for biomedical applications.

## Description

Calcium phosphate-based transfection is a versatile method for introducing foreign genetic material into eukaryotic cells (Goetze et al., 2004; Guo et al., 2017; Kwon & Firestein, 2013). This approach forms a calcium phosphate-DNA coprecipitate, effectively delivered to the target cells (Jordan & Wurm, 2004). Widely recognized for its universality, cost-effectiveness, and minimal cellular perturbation, this technique proves invaluable in molecular biology and genetic engineering.

However, challenges persist, including variable efficiency among different cell types, potential cytotoxicity, and limitations in payload capacity (Lo et al., 2019). Although the method is applicable across diverse cell types and cost-effective compared to alternatives, its invasiveness varies and may pose cytotoxic risks, particularly in specific cell types (Carballo-Pedrares et al., 2023; Chong et al., 2021). Scaling up for industrial applications proves challenging, and complexities arise with more extensive or multiple plasmids. Understanding these nuances is essential for researchers employing calcium phosphate-based transfection, ensuring optimal experimental design and outcomes.

Human splenic fibroblast cells serve critical functions within the spleen, contributing to structural and immunological aspects. These cells are primary contributors to the structural integrity of the spleen through the synthesis of collagen, a vital component of the extracellular matrix (ECM) (Lokmic et al., 2008). Splenic fibroblasts actively participate in immune responses by producing cytokines, chemokines and interleukins, influencing inflammation, immune cell recruitment, and immunomodulation (Correa-Gallegos et al., 2021; Zhao et al., 2021).

In wound healing and tissue repair, splenic fibroblasts undergo fibroplasia, which is pivotal in regenerating damaged tissues (de Oliveira & Wilson, 2020). Furthermore, these cells contribute to the dynamic regulation of the ECM, influencing the spleen’s mechanical properties and overall functionality (Bellomo et al., 2021).

Splenic fibroblasts provide essential support for immune cells by creating a framework within the spleen, facilitating the organization and functioning of lymphocytes and other immune cells (Crane et al., 2021; Lewis et al., 2019; Mebius & Kraal, 2005; Noble et al., 2018). In pathological conditions, changes in the behaviour of splenic fibroblasts may impact immune responses, making their study crucial for understanding the role of the spleen in various diseases (Lewis et al., 2019).

These cells may also play a role in pathogen recognition and response, contributing to the spleen’s intricate immune defence mechanisms (Lewis et al., 2019). As valuable experimental models, human splenic fibroblasts provide insights into cell behaviour, immunological processes, and responses to external stimuli, advancing biomedical research (Pezoldt et al., 2021). Moreover, their involvement in immune regulation and tissue repair positions splenic fibroblasts as potential therapeutic targets in conditions involving immune dysregulation or tissue damage. Comprehending the diverse roles played by human splenic fibroblast cells is crucial for unravelling the intricacies of spleen function and its impact on immunity.

In human splenic fibroblasts, transfection is essential for understanding gene expression modulation within a physiologically relevant setting. Challenges specific to human splenic fibroblasts include their primary cell nature, cell-type specificity, and cultural variability among donor-derived fibroblast cultures. Addressing these difficulties is crucial for achieving reliable outcomes in experiments and advancing basic research and potential therapeutic applications.

In culture, normal primary human splenic fibroblast cells exhibit a distinctive spindle-shaped morphology characterized by elongated, fibrous structures extending from the cell body (Figure 1: A1-A3) (Gauthier et al., 2023). These cells adhere closely to the culture substrate, forming a monolayer with a flattened appearance. The nuclei are generally oval or elongated, centrally located within the cell, and surrounded by a well-defined cytoplasm (Figure 1: A1-A3). This cellular architecture reflects a dynamic interplay between cell-substrate interactions and intrinsic cellular processes, which is crucial for maintaining a healthy and functional fibroblast population. The spindle shape indicates their mesenchymal origin, which is essential in supporting tissue structure and function within the spleen (Dominici et al., 2006). Understanding this characteristic morphology is paramount when investigating the behaviour and function of normal primary human splenic fibroblast cells in a cultured environment.

**Figure 1:**
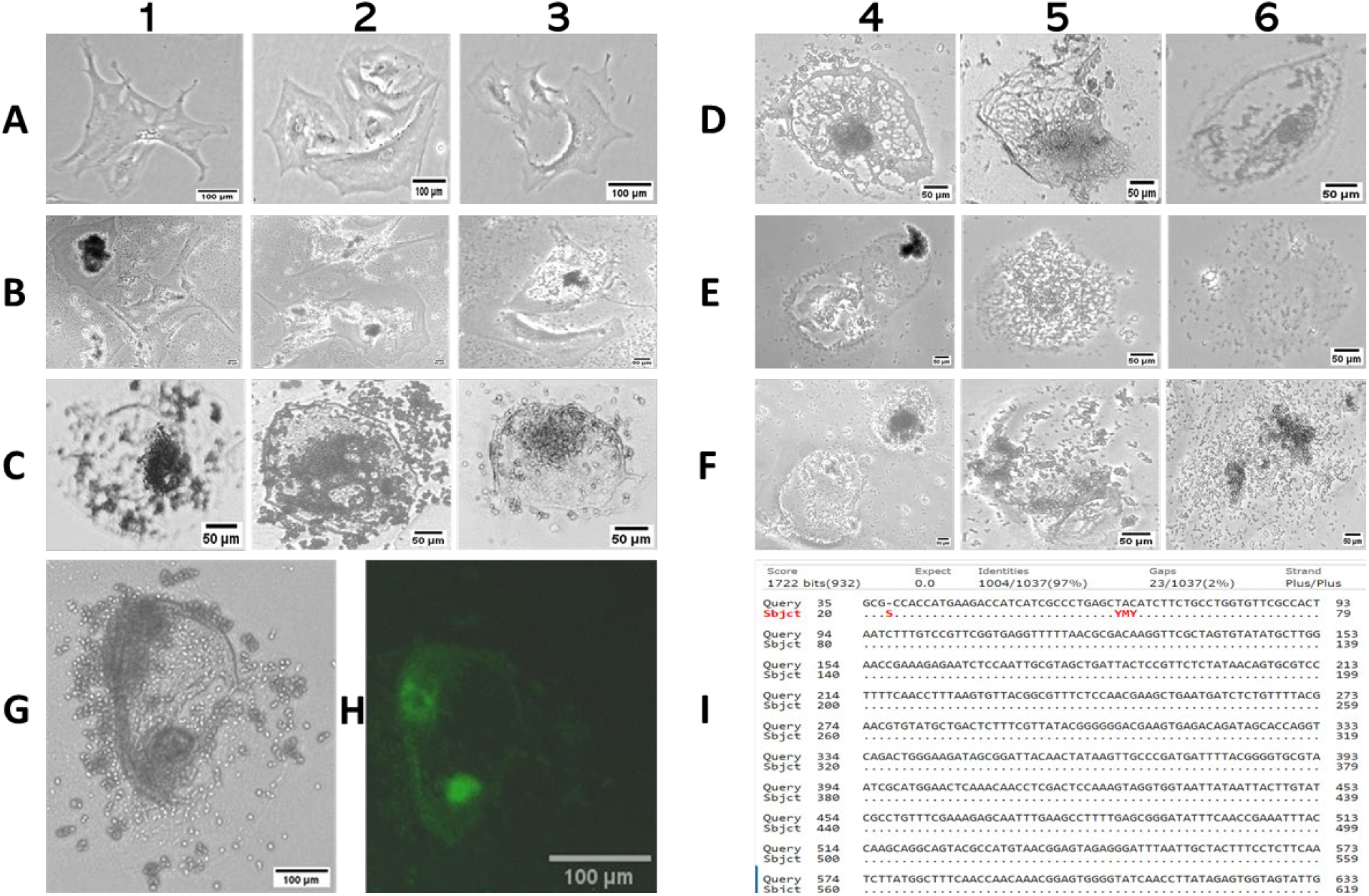
The micrographs presented in Figure 1 depict human splenic fibroblast cells in both control conditions (Figure 1: A1-A3) and under transfection with pcDNA3-SARS-CoV-2-S-RBD-sfGFP using Promega’s calcium phosphate reagent (Figure 1: B1-F3) at varying time points (4, 8, 24, 48, and 72 hours). These images provide a detailed examination of the dynamic impact of the plasmid and transfection reagent on the morphology and behaviour of HSFC over time. The upper-left panel (Figure 1: A1-A3) serves as a reference, showcasing the characteristic growth patterns of normal HSFC without the transfection reagent. In contrast, subsequent panels (Figure 1: B1-F3) vividly illustrate the consistent influence of the transfection process on altering fibroblast shapes and behaviours throughout the experimental timeline. Noteworthy snapshots at 8 hours post-transfection (Figure 1: G and H) precisely capture individual cells, revealing dynamic changes induced by vesicles. Figure 1: (I) complements these micrographs by presenting the sequencing result of the plasmid utilized during transfection. These comprehensive observations extend beyond morphological shifts, emphasizing the profound impact of the transfection process on the cellular microenvironment. This study contributes valuable data to the broader scientific discourse, shedding light on the intricate interplay between transfection processes and cellular responses. The findings hint at potential avenues for understanding how experimental interventions intricately modulate cell shape, growth, and behaviour, thereby enriching our understanding of transfection methodologies in biomedical research.

Upon treatment with calcium-phosphate-DNA, there was a significant alteration in the cell morphology, leading to a reduction in the spindle and elongated shape. Four hours post-transcription conditions revealed an observable beginning shift to an oval cell’s shape (Figure 1: B1-B3). This change persisted at the subsequent stage, 8 hours post-transfection, where the cells rounded up. This physiological regression in cell shape suggests a reaction to the transfection reagent and plasmid. Notably, at 8 hours post-transfection, numerous vesicles were observed around the nuclei in the cytoplasm and the extracellular space (Figure 1: C1-C3, G, and H). Initial assumptions regarding these blisters resulting from the reagent, through exocytosis-phagocytosis due to calcium phosphate precipitating the DNA, were deemed invalid. Calcium phosphate transfection induces changes in cellular calcium levels, prompting cellular responses such as vesicle formation related to calcium homeostasis or stress responses. The transfection process may also trigger the release of extracellular vesicles, including exosomes, responding to the stress introduced by the transfection reagents. Introducing foreign DNA or transfection reagents could stimulate an innate immune response in the fibroblasts, releasing vesicles involved in immune signaling or response. The specific observation of vesicles at this stage, distinct from other conditions, requires further investigation to understand better the cells’ reaction and the nature of the vesicles at this particular time.

Subsequent time points in other cell conditions illustrate a progression towards cellular disintegration, culminating in cells falling apart by 72 hours post-transfection (Figure 1: E1-F3). The study identifies an 8-hour critical window for successful transfection, beyond which temporal toxicity leads to cell disintegration. Moreover, the unique spindle-shaped morphology of splenic fibroblasts undergoes alterations post-transfection, with vesicle formation suggesting cellular responses related to calcium homeostasis or stress responses. Further investigation into vesicle nature and immune responses is warranted, offering comprehensive insights for broader biomedical research applications and potential therapies.

## Methods

### Cell Preparation

Cell culture was conducted in T75 flasks (GeneBio System, ON, Canada) laid the foundation for rigorous experimentation. A week before transfection, the old medium was aspirated and discarded. The adherent cell layer underwent four rinses with 40 ml sterile 1X PBS lacking calcium and magnesium (Invitrogen). Using a sterile pipette facilitated the dislodging of loosely attached cells and the elimination of debris. The rinse solution was discarded, and the cells were treated with 3.0 ml of 0.05% Trypsin-EDTA solution (Cell Biologics, Catalog No. 6915) for 8 minutes, facilitating complete cell detachment through gentle tapping of the flask. Subsequently, 10 ml of Cell Biologics’ Cell Culture Medium, supplemented with 10% FBS, neutralized the trypsin-EDTA reaction. Following cell counting using ImageJ, 10^4^ cells were seeded into each well of a 24-well plate. The plate was then incubated in a humidified, 5% CO2 incubator at 37°C. Daily monitoring of cell morphology persisted until reaching 50% confluence, ensuring optimal conditions for subsequent experimental procedures. This systematic approach establishes a robust cellular environment for transfection studies, emphasizing meticulous attention to cell detachment, seeding density, and culture maintenance for accurate and reproducible results.

### Plasmid

The pcDNA3-SARS-CoV-2-S-RBD-sfGFP plasmid, generously provided by Dr. Erik Procko from Illinois State University (Addgene plasmid #141184), is a critical component of this study. The plasmid has a total size of 6736 base pairs (bp), including a vector backbone size of 5335 bp and an insert size of 1401 bp representing the spike protein. The cloning methodology involved a 5′ cloning site at NheI (undisturbed) and a 3′ cloning site at XhoI (maintained). The 5′ sequencing primer used was CMV-fwd, while the 3′ sequencing primer employed was BGH-rev. The resulting cloning-grade DNA was validated for utility in PCR, cloning reactions, or transformation into E. coli. However, further considerations were required to optimize purity and quantity for direct transfections in subsequent experimental procedures.

### Transformation and bacterial culture

The pcDNA3-SARS-CoV-2-S-RBD-sfGFP plasmid was introduced into Trans5alpha Chemically Competent cells through a meticulous transformation. Briefly, Trans5alpha Chemically Competent Cells were prepared using a standard calcium chloride treatment protocol. The plasmid DNA was added to the competent cells with utmost precision. The transformation mixture underwent a controlled heat shock followed by rapid cooling, inducing the uptake of the plasmid by the competent cells. After the heat shock, a recovery period in a nutrient-rich medium allowed for cell recuperation and plasmid expression. The transformed cells were subsequently plated onto selective agar containing an appropriate antibiotic for plasmid selection. Following successful transformation, colonies were meticulously chosen for cultivation, employing ampicillin as a selective agent on agarose gel and LB broth. Subsequent mini and midi preps, guided by Qiagen instructions, were carried out. Verifying the pcDNA3-SARS-CoV-2-S-RBD-sfGFP plasmid’s presence involved analytical techniques, including 0.8% agarose gel electrophoresis. Additionally, specific regions of the plasmid were digested and linearized using restriction enzymes (BamHI, NheI, and XbaI). Sequencing, utilizing the sequence 5’ – TAA TAC GAC TCA CTA TAG GG – 3’ was a gift from KPU Applied Genomics Centre (AGC) directed against the T7 present in the plasmid, ensured the establishment of a stable and genetically modified Trans5alpha Chemically Competent Cell harbouring our plasmid construct. This methodological approach adhered to established best practices for efficient and reproducible plasmid transformation into bacterial cells.

### Transfection and observation of cell morphology

The system components were thawed to room temperature. Each component underwent thorough mixing by vortexing for optimal homogeneity. Three hours preceding transfection, the cell culture medium was replaced with a fresh growth medium. DNA and 2X HBS solutions were meticulously prepared for each transfection in separate sterile tubes. DNA and water were combined in the first tube, followed by adding CaCl_2_ with subsequent mixing. The specified volume of 2X HBS was added to the second tube. Working within a sterile tissue culture hood, the tube containing the 2X HBS solution was gently vortexed, adjusting the speed for safe vortexing without the cap and allowing the prepared DNA solution to be added. Simultaneously, the prepared DNA solution from series of tubes 1 was introduced dropwise into the 2X HBS in series of tubes 2, ensuring constant vortexing. The solution exhibited a slightly opaque appearance upon completion, indicating the fine calcium phosphate-DNA coprecipitate formation. Incubation at room temperature for 30 minutes followed. Before addition to cells, the transfection solution was vortexed again. The solution containing the CaCl_2_-DNA residue was added dropwise to the cells at various plate locations. Gentle swirling ensured even distribution, preventing localized acidification. The plates were then returned to a humidified 5% CO^2^ at 37°C for further incubation. On day 3 post-transfection, data were collected from cells (Figure 1: A1-H). The transfection state, lasting 8 hours, achieved an 85-95% transfection rate, indicating improved conditions and timing for this cell cohort. Micrographs were captured using Leica MC120 HD and EVOS FL Color (Life Technologies). This data contributes to our understanding of cell behaviour and responses in the specified post-transfection timeframe, facilitated by advanced microscopy techniques for comprehensive visualization.

## Acknowledgments

We sincerely thank Dr. Monica De Boer at Kwantlen Polytechnic University (KPU)-Surrey campus for generously providing ProFection (E1200, WI, Promega). Ensuring the non-infectious nature of pcDNA3-SARS-CoV-2-S-RBD-sfGFP was of utmost importance, and we are indebted to Dr. Eric Procko from Illinois State University for his expertise in confirming the plasmid’s safety. His validation enables the secure handling of the plasmid at biosafety levels 1 or 2, effectively preventing any risk of SARS-CoV-2 virus transmission. Special thanks to the KPU Applied Genomic Center (AGC) team’s generous gift of the T7 primer, a crucial component in generating the sequence of our plasmids. We thank Dr. Layne Myhre and Sean Conway (KPU) for their unwavering support and encouragement at various stages. Their significant contributions have greatly influenced the success of this manuscript.

## Author Contributions

Ashpreet Kaur: Data curation, Formal analysis, Contributed to the investigation, methodology, software, and visualization.

Victoria A. Assogba: Contributed to the investigation, Data curation, Formal analysis, Methodology, Writing - review, and editing.

Jovy M. G. Assogba: Data curation, Formal analysis, Writing – review and editing.

Barnabe D. Assogba: Conceptualization, Data curation, Formal analysis, Funding acquisition, Investigation, Methodology, Project administration, Supervision, Writing - original draft, Writing – review and editing.

